# CYFIP1 overexpression amplifies IL-6/STAT3 and IFN-gamma/STAT1 signaling: potential implications for neuroinflammation and autism spectrum disorder

**DOI:** 10.1101/2025.10.23.683866

**Authors:** Emily-Rose Martin, Josan G. Martin, Mark A. Russell, Asami Oguro-Ando

## Abstract

Autism spectrum disorder (ASD) encompasses a group of neurodevelopmental disorders characterised by impaired social interaction, delayed language development, and repetitive or restrictive behaviours. While both genetic and environmental factors contribute to ASD, the specific molecular mechanisms underlying these interactions remain unclear. In a recent preprint, we showed that cytokine signaling may be dysregulated in chromosome 15q-duplication syndrome (Dup(15q)), one of the most common syndromic forms of ASD. Specifically, Dup(15q) induced pluripotent stem cells (iPSC)-derived neurons exhibit an amplified Signal Transducer and Activator of Transcription-3 (STAT3) response to Interleukin-6 (IL-6), a proinflammatory cytokine often upregulated in ASD. To identify which gene in the 15q region may be responsible for modifying cytokine signalling responses in Dup(15q), we focus on the *Cytoplasmic FMRP-Interacting Protein 1 (CYFIP1*) gene, as increased *CYFIP1* dosage in Dup(15q) is associated with increased severity of neurobehavioral symptoms in individuals with Dup(15q) and *CYFIP1* dysregulation has been linked to altered expression of various genes involved in immunoregulatory signalling pathways. Here, we explore the effects of *CYFIP1* overexpression (*CYFIP1*-OE) on cytokine signaling, demonstrating that *CYFIP1*-OE in HEK293 cells modifies the expression and activity of several cytokine signaling-related transcription factors, including STAT3 and STAT1. Additionally, we use SH-SY5Y neuroblastoma cells to assess neuron-related phenotypes, showing that *CYFIP1*-OE alters IL-6-induced neurite outgrowth. These findings provide novel insights into how CYFIP1 may contribute to dysregulated cytokine responses in ASD, advancing our understanding of the molecular mechanisms that underlie neuroinflammatory processes in this disorder.

## Background

Autism spectrum disorder (ASD) is a heterogeneous group of neurodevelopmental conditions in which both genetic variation and environmental exposures contribute to pathogenesis (1). Due to the highly heterogeneous nature of the disorder, both diagnosis and treatment remain challenging. Genetic factors, including both rare and common variants, are implicated in ASD, with over 1,000 genes identified as potentially contributing to its onset and development (2). In addition to genetic predisposition, environmental risk factors, particularly maternal immune activation (MIA) and neuroinflammation, have been recognised as key contributors to ASD pathogenesis (3,4). Therefore, it is likely that the interaction between an individual’s genetic background and specific environmental exposures plays a crucial role in triggering the pathogenesis underlying ASD (3). Consequently, elucidating the molecular mechanisms by which ASD-associated genes can alter neurodevelopmental processes in response to environmental stressors is critical for advancing our understanding of the disorder.

*Janus Kinase and Microtubule-Interacting Protein 1* (*JAKMIP1*) was previously identified as a potential molecular link between syndromic forms of ASD and neuroinflammation (5). *JAKMIP1* expression was commonly dysregulated in Fragile X syndrome (FXS) and chromosome 15q-duplication syndrome (Dup(15q)), both genetic disorders often comorbid with ASD (5). Furthermore, we discovered a role for JAKMIP1 in regulating Interleukin-6 (IL-6)-induced Signal Transducer and Activator of Transcription-3 (STAT3) signaling in neuronal cells, whereby deficiency of JAKMIP1 in SH-SY5Y neuroblastoma cells impairs STAT3 activation and neuritogenesis in response to IL-6 stimulation (5). These findings suggest that dysregulation of *JAKMIP1* in ASD may alter the ability of neurons to respond to inflammatory cytokines, potentially leading to aberrations in neuronal morphology.

To investigate whether *JAKMIP1* dysregulation in syndromic ASD could modify how cells respond to cytokines, we previously assessed IL-6-induced STAT3 signalling in a Dup(15q) induced pluripotent stem cell (iPSC)-derived neuronal model (6), as Dup(15q) is one of the most common forms of syndromic ASD, accounting for around 1% of ASD cases (7). Compared to a control iPSC line, we observed that *JAKMIP1* expression fluctuates in Dup(15q) iPSC and iPSC-derived cortical neurons (iNeurons), being downregulated in Dup(15q) iPSC and upregulated in iNeurons. Despite these opposing trends, we found that *STAT3* expression was consistently reduced in both the Dup(15q) iPSCs and iNeurons, indicating that Dup15q cells would likely display altered responses to IL-6. Curiously, though *STAT3* expression was reduced, IL-6-induced STAT3 activity was significantly enhanced in both Dup(15q) iPSCs and iNeurons. These findings suggest that unlike in the SH-SY5Y cells, JAKMIP1 is likely not regulating *STAT3* expression or activity in Dup(15q) and instead, another gene in the chromosome 15q region exerts stronger influence on STAT3 activity.

Several genes are encoded within the 15q region of chromosome 15, however, a gene of particular interest in relation to ASD and cytokine signalling is *Cytoplasmic FMRP-Interacting Protein 1* (*CYFIP1*), as studies have demonstrated that duplications (or even deletions) encompassing *CYFIP1* increase the severity of neurobehavioural phenotypes (8), and that *CYFIP1* expression is upregulated in the brains of individuals with Dup(15q) and ASD (9). Although CYFIP1 is most known for its roles in repressing neuronal protein translation as part of Fragile X Messenger Ribonucleoprotein (FMRP)-containing ribonucleoprotein complexes (10) and regulating actin polymerisation as a core component of the WAVE regulatory complex (WRC) (11), there is evidence to suggest that *CYFIP1* dysregulation may alter the expression of various genes involved in immunoregulatory signalling pathways (12). For example, a recent study profiling transcriptomic changes in Dup(15q) stem cell-derived neurons with upregulated *CYFIP1* expression suggested that increased dosage of chromosome 15q11.2 genes could alter the expression gene targets of SMAD Family Member 3 (SMAD3), Nuclear Factor Kappa B (NF-κB) or Interferon-Stimulated Response Elements (ISRE) (12). Moreover, another study identified *CYFIP1* as a potential hub gene that may coordinate Transforming Growth Factor Beta 1 (TGF-β1)/SMAD3 signalling (13). Therefore, we hypothesised that CYFIP1 may be responsible for the observed enhanced IL-6-induced activity in Dup(15q) iPSCs and iNeurons (6).

This paper therefore explores the effects of increased *CYFIP1* expression (modelled by *CYFIP1*-overexpression (OE)) on cytokine signalling. We provide the first evidence to directly support a novel role for CYFIP1 in modulating cytokine signalling, demonstrating that *CYFIP1*-OE in HEK293 cells alters the expression and activity of cytokine signalling-related transcription factors, *STAT1* and *STAT3*. Taking this further, we show how alterations in the STAT3 signalling pathway due to *CYFIP1*-OE affects IL-6-induced neurite outgrowth in human SH-SY5Y neuroblastoma cells, illustrating that these findings continue to be relevant in a more neuronal-relevant model.

## Methods

### HEK293 and SH-SY5Y cell culture and treatment

Human embryonic kidney (HEK)-293 cells (#CRL-1573™, *American Type Culture Collection*) and human neuroblastoma cells (#CRL-2266™, *American Type Culture Collection*) were thawed from frozen stocks stored in 10% dimethyl sulfoxide (#D2650-5X5ML, *Sigma-Aldrich*®) at −80°C. HEK293 cells and SH-SY5Y cells were maintained in 10-cm diameter Petri dishes (#P7612-360EA, *Sigma-Aldrich*®) with 10 mL Dulbecco’s modified eagle medium/nutrient mixture F-12 with GlutaMAX (referred to henceforth as DMEM/F-12) (#11524436, *Fisher Scientific*™) supplemented with 10% foetal bovine serum (FBS) (#11550356, *Fisher Scientific*™), and incubated at 37°C, 5% CO_2_, 95% humidity. Cells were passaged at 70-80% confluency using 3 mL of pre-warmed TrypLE™ (#10718463, *Fisher Scientific*™) for 5 min and TrypLE™ was inactivated by dilution in an equal volume of DMEM/F-12 + 10% FBS.

Cells were stimulated with 20 ng/mL IL-6 (#7270IL-025, *R&D Systems*) or 20 ng/mL Interferon-γ (IFN-γ) (#285-IF-100, *R&D Systems*) diluted in DMEM/F-12 + FBS (or in DMEM/F-12 supplemented with Retinoic Acid (RA) and Brain-Derived Neurotrophic Factor (BDNF) for neurite tracing experiments). For all experiments using IL-6 or IFN-γ, control cells were used which were treated with corresponding media not containing IL-6 or IFN-γ. Cells were treated with IL-6 or IFN-γ for different amounts of time depending on the purpose of the experiment. To assess STAT3 activation by Western blotting for phosphorylated STAT3 (P-STAT3^Y705^), cells were treated with IL-6 for 30 minutes. To assess STAT3 or STAT1 transcriptional activity by Dual-Luciferase® Reporter (DLR™) assays, cells were treated with IL-6 or IFN-γ for 18 hours post-transfection. To measure mRNA expression of STAT3 or STAT1 responsive genes, cells were treated with IL-6 or IFN-γ for 4 hours. Finally, to assess IL-6-induced neurite outgrowth, cells were treated with IL-6 over a 48-hour period of differentiation.

### HEK293 cell transfection

24 hours prior to transfection, 300,000 HEK293 cells were seeded into Nunc cell-culture treated 6-well plates (#10469282, *Fisher Scientific*™) in 3 mL DMEM/F-12 + 10% FBS. The following day, HEK293 cells were transfected with either an RFP-T2A-CYFIP1 plasmid to induce *CYFIP1*-overexpression (these cells are referred to as ‘*CYFIP1*-OE cells), or the backbone RFP-T2A plasmid as a control (these cells are referred to as ‘control cells’) using Lipofectamine™ LTX Reagent (#15338100, *Fisher Scientific*™) following manufacturer instructions (i.e., final amount of 2.5 μg of plasmid and 5 μL of Lipofectamine™ LTX per well). Media was replaced the following day. All experiments were performed 72 hours post-transfection.

### SH-SY5Y cell transfection

2,000,000 SH-SY5Y cells were transfected by Nucleofection™ (#V4XC-2012, *Lonza Bioscience*) following manufacturer instructions with 5 μg of either a mixture of the RFP-T2A backbone vector and pCAβ-YFP (encoding yellow fluorescent protein (YFP)) in a 1:1 ratio as the control condition; or a mixture of RFP-T2A-CYFIP1 and pCAβ-YFP (4:1 ratio) to assess the effects of *CYFIP1*-OE on neuritogenesis. A pCAβ-YFP plasmid was co-transfected to enable better visualisation of the cells for neurite tracing as the RFP encoded by the *CYFIP1*-OE plasmid photo-bleaches quickly. Media was replaced the following day.

### Protein extraction and Western blotting

72 hours post-transfection, cells were washed with ice-cold 1x phosphate buffered saline (1xPBS) (#P4417-50TAB, Sigma-Aldrich®) then collected with a cell scraper in 300 μL of ice-cold RIPA lysis buffer (150 mM NaCl, 50 mM Tris/Cl pH = 8.0, 0.5% Sodium Deoxycholate, 0.1% SDS, 1% Triton X-100) supplemented with 1 mM PMSF, 1% Phosphatase Inhibitor Cocktail 2 (#P5726-1ML, Sigma Aldrich®), 1% Phosphatase Inhibitor Cocktail 3 (#P0044-1ML, Sigma Aldrich®) and 1% Complete Protease Inhibitor Cocktail (#P8340-1ML, Sigma Aldrich®). Cell lysates were incubated on ice for 30 minutes, cleared by centrifugation at 17,000xg for 15 minutes at 4°C, then supernatants collected. Bicinchoninic Acid assays (#10678484, *Thermo Scientific*) were performed to estimate protein concentrations and 50 μg of total protein was diluted with Laemmli buffer (final concentration of 60 mM Tris/Cl pH = 6.8, 10% Glycerol, 2% SDS, 5% 2-Mercaptoethanol, 0.02% Bromophenol Blue), then boiled for 5 minutes at 95°C. 50 μg of total protein for each sample was loaded into 4–15% precast polyacrylamide gels (#4561085, *Bio-Rad Laboratories*) alongside a pre-stained protein ladder (#PL00001, *Proteintech Group*). Electrophoresis was performed in the Mini-PROTEAN® Tetra Vertical Electrophoresis Cell (#1658004, *Bio-Rad Laboratories*) with 1×SDS-PAGE running buffer (2.5 mM tris-base; 19 mM glycine (#10070150, *Fisher Chemical*™); 0.01% SDS; pH = 8.3 in ddH_2_O). Proteins were then transferred onto methanol-activated PVDF membranes (#IPVH304F0, *Millipore*) by a wet transfer method in the Mini-PROTEAN® Tetra Vertical Electrophoresis Cell. PVDF membranes were then blocked for at least 1 hour at room temperature with 5 mL of 5% skim milk in 1x tris-buffered saline (TBS; 2 mM Tris-Base; 15 mM NaCl; pH = 7.6 in ddH_2_O) supplemented with 0.1% Tween® 20 (#10485733, *Fisher BioReagents*™) (1xTBS-T). After blocking, membranes were sequentially incubated with 5 mL of primary antibody (gentle rocking for 2-3 hours at room temperature or overnight at 4°C), then secondary antibody (gentle rocking for 30 minutes-1 hour at room temperature). Primary and secondary antibodies were diluted in 5% skim milk in 1x TBS-T (or 5% BSA in 1x TBS-T for anti-P-STAT3^Y705^). The dilution factors and specific antibodies used are listed in **Table 1**. Three 5-minute washes with 1x TBS-T were performed between antibody incubations. GAPDH was used as a loading control to normalise expression of target proteins. Proteins were visualised using the Odyssey® CLx Imager (*LI-COR Biosciences*) and the Image Studio software (Version 5.2; *LI-COR Biosciences*) was used to perform densitometric analyses to compare the abundance of target proteins between samples.

**Table 1:** Antibodies used in this study.

| Catalogue Number | Supplier | Host Species | Target | Dilution | Conjugated Fluorophore |
| --- | --- | --- | --- | --- | --- |
| ab156016 | *Abcam* | Rabbit | CYFIP1 | 1:1000 |  |
| sc-47724 | *Santa Cruz Biotechnology* | Mouse | GAPDH | 1:1000 |  |
| 13846-1-AP | *Proteintech* | Rabbit | JAKMIP1 | 1:1000 |  |
| 9139S | *Cell Signaling Technology* | Mouse | STAT3 | 1:1000 |  |
| 9131l | *Cell Signaling Technology* | Rabbit | P-STAT3<sup>Y705</sup> | 1:1000 |  |
| 35519 | *Invitrogen* | Goat | Mouse | 1:5000 | DyLight™ 680 |
| SA5-10036 | *Invitrogen* | Goat | Rabbit | 1:5000 | DyLight™ 800 |

### RNA extraction, cDNA conversion and qRT-PCR

72 hours post-transfection, total RNA was isolated and purified using the Direct-zol™ RNA Miniprep kit following manufacturer instructions (#R2051, *Zymo Research*). 500 ng of RNA was converted to cDNA using PrimeScript™ RT reagent kit (#RR037A, *Takara Bio Europe*) following manufacturer instructions. qRT-PCR was then performed using HOT FIREPol® EvaGreen® qPCR Master Mix with ROX (#01-02-00500, *Solis BioDyne*) with the QuantStudio 12K Flex qPCR machine (*Thermo Fisher Scientific*™). For primer sequences, see **Table 2**. mRNA expression levels were standardized against two reference genes, *Glyceraldehyde-3-Phosphate Dehydrogenase* (*GAPDH*) and *RNA Polymerase II Subunit A* (*POLR2A*) using the Pfaffl method (14) and normalised to the control cells.

**Table 2:**
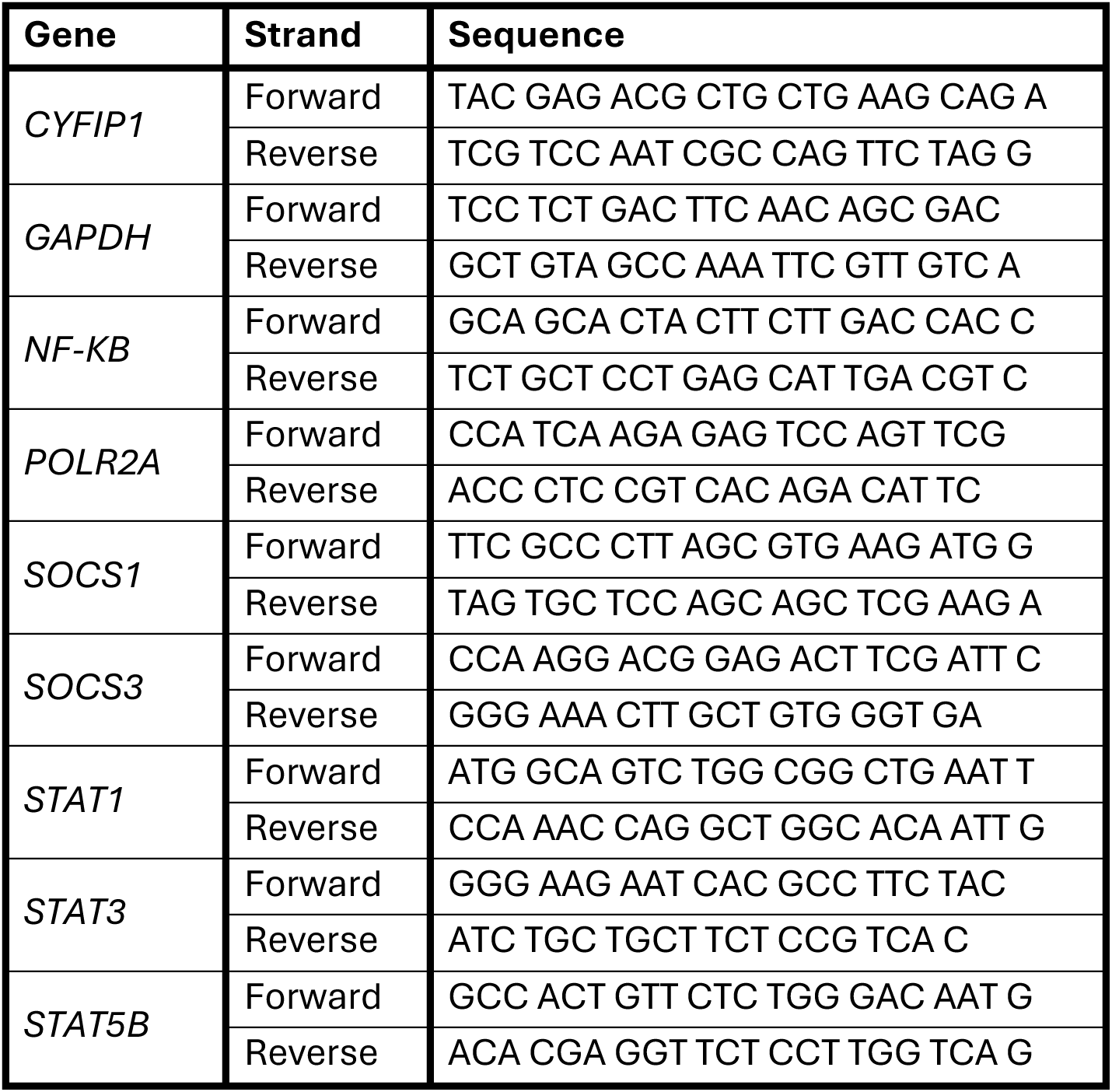
Primers used in this study.

### Dual-luciferase reporter assays

50,000 HEK293 cells were seeded into 24-well plates and incubated overnight. The following day, cells were transfected with 2 μL of STAT3 Cignal Reporter plasmid (#336841, GeneGlobe ID: CCS-9028L, *Qiagen*) or 2 μL of STAT1 (GAS) Cignal Reporter plasmid (#336841, GeneGlobe ID: CCS-009L, *Qiagen*) using Lipofectamine™ LTX as previously described. Four hours post-transfection, cells were treated with DMEM/F-12 +10% FBS with or without IL-6- or IFN-γ and incubated overnight. 18 hours later, Firefly and Renilla luciferase activities were measured using the Dual-Luciferase® Reporter (DLR™) Assay System (#E1960, *Promega*) following manufacturer instructions. Luminescence was measured in white-bottomed LUMITRAC™ 96-well plates (#655074, *Greiner Bio-One*) using the PHERAstar FS microplate reader (*BMG LABTECH*).

### SH-SY5Y diSerentiation and neurite outgrowth analysis

Sterile 13-mm-diameter glass coverslips (#631-1578, *VWR*®) were placed into 24-well plates and coated with 300 μL of 20 μg/mL poly-D-Lysine (PDL; #354210, *Corning*®) by overnight incubation at 37°C. 24 hours post-transfection, 30,000 SH-SY5Y cells were seeded onto PDL-coated coverslips in DMEM/F-12 + 10% FBS. The following day, media was replaced with 20 ng/mL BDNF and 10 μM RA in DMEM/F-12 (without FBS). 48 hours later, SH-SY5Y cells were fixed with 4% paraformaldehyde (PFA), and nuclei were counterstained with 4′,6-diamidino-2-phenylindole (DAPI) (#D9542, *Sigma-Aldrich*®) for 15 minutes. Coverslips were mounted onto glass microscope slides with ProLong™ Diamond Antifade Mountant (#15468070, *Fisher Scientific*™) and slides were imaged on the inverted DMi8 widefield microscope (*Leica Microsystems*).

All image analysis was performed manually using the *FIJI* software (*ImageJ*; Version 2.9.0; (15)). Prior to image analysis, a scale bar was used to set a pixel to μm ratio to each image, after which the “Segmented Line” tool was used to measure the distance from the edge of the nucleus to the furthest end of each neurite. The length of the longest neurite produced by a single cell (referred to as longest neurite length (LNL); a measure of neurite extension) and the sum length of all neurites produced by a single cell (referred to as total neurite length (TNL); a measure of neuritogenesis) were measured.

### Statistical analysis

Statistical analyses were performed using GraphPad Prism (GraphPad Prism Version 9.0.0 for Windows, GraphPad Software, San Diego, California USA, www.graphpad.com). Unpaired Student’s t-Tests were used to assess statistical differences in measurements between two groups (e.g., *CYFIP1*-OE against control). When investigating the effects of and interactions between two independent variables (e.g., the effect of both *CYFIP1*-OE and IL-6 treatment on STAT3 transcriptional activity), two-way analysis of variance (ANOVA) were performed. Appropriate post hoc tests were also performed to account for multiple testing. Data are expressed as mean values ± standard error of the mean (SEM). P-values < 0.05 were considered statistically significant.

## Results

### Modelling CYFIP1-overexpression

To model increased *CYFIP1* expression, we utilised plasmid vectors encoding an RFP-T2A-CYFIP1 construct (to induce *CYFIP1*-OE) or RFP-T2A as a control, which we transfected into HEK293 cells, referred to as *CYFIP1*-OE and control cells respectively (**Figures 1A and 1B**). We then confirmed increased *CYFIP1* mRNA expression using qRT-PCR (**Figure 1C**) and protein expression using Western blotting (**Figures 1D and E**).

**Figure 1.**
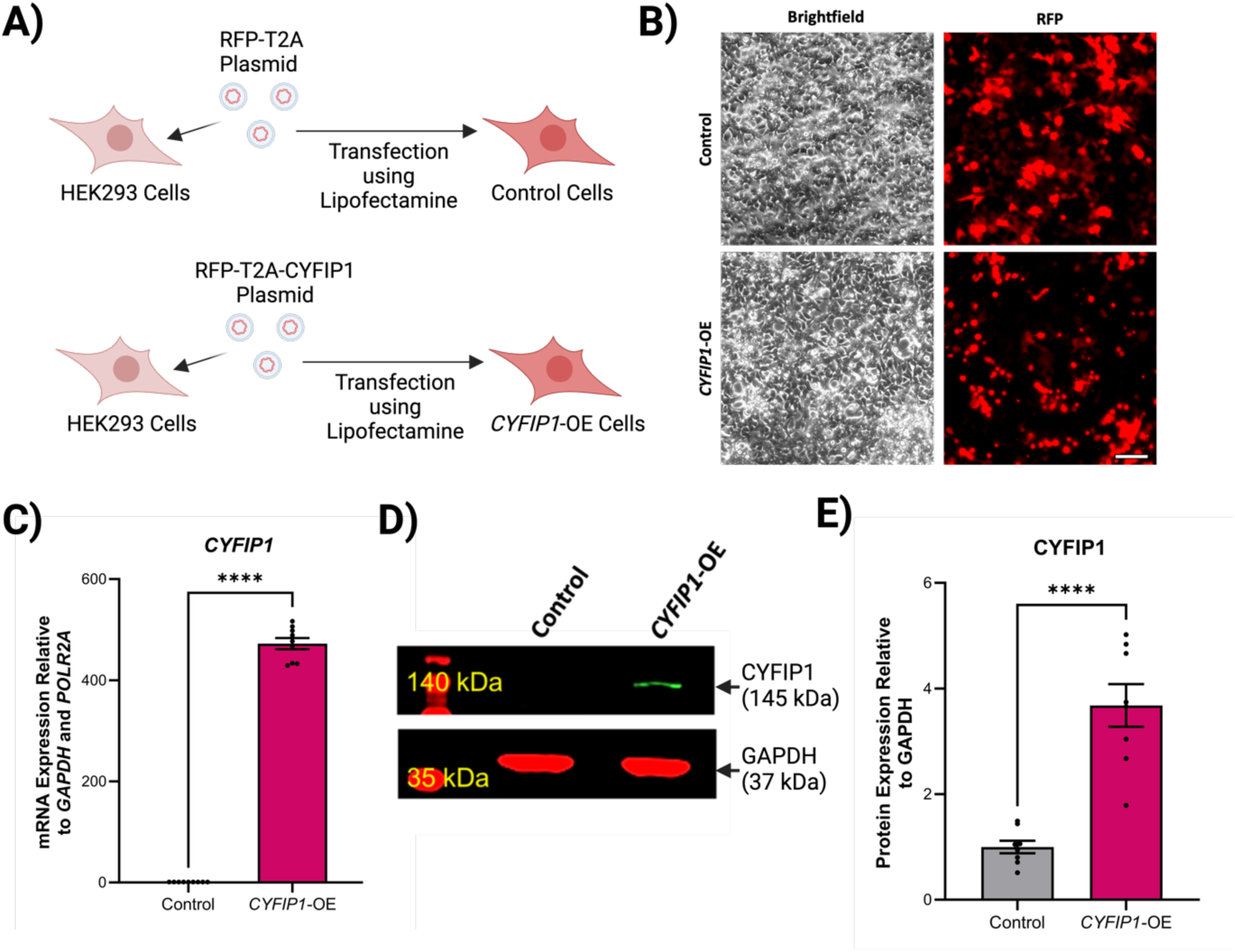
| Modelling CYFIP1 overexpression in HEK293 cells. **A)** HEK293 cells were transfected with either RFP-T2A plasmid as a control (referred to as Control), or RFP-T2A-CYFIP1 to induce CYFIP1-overexpression (OE) (referred to as CYFIP1-OE). Created with BioRender.com. **B)** Control and CYFIP1-OE cells were imaged 72 hours post-transfection live on the Leica DMi8 widefield microscope. Scale bar = 100 μm. **C)** Relative mRNA expression of CYFIP1 in Control and CYFIP1-OE cells 72 hours post-transfection measured by qRT-PCR. **D)** Increased CYFIP1 protein expression in CYFIP1-OE cells 72 hours post-transfection confirmed via Western blotting. **E)** Densitometric analysis of total CYFIP1 protein expression relative to the GAPDH loading control. All values are normalised to the Control cells. For all experiments, values are presented as mean ± SEM of N = 3 independent experiments, 2-3 technical replicates each. Statistical significance against Control was determined using Unpaired Student’s T-Tests; ****P< 0.0001.

### *CYFIP1*-OE leads to reduced JAKMIP1 expression

It has been previously demonstrated that *JAKMIP1* expression is reduced following *CYFIP1*-OE in human neuroblastoma cells (5). Therefore, to confirm whether this is the case in HEK293 cells, we measured *JAKMIP1* expression in *CYFIP1*-OE HEK293 cells by qRT-PCR and Western blotting. Agreeing with Nishimura et al., we found that *JAKMIP1* mRNA (Figure 2A) and protein expression (Figure 2B and C) is significantly reduced in *CYFIP1*-OE HEK293 cells.

**Figure 2.**
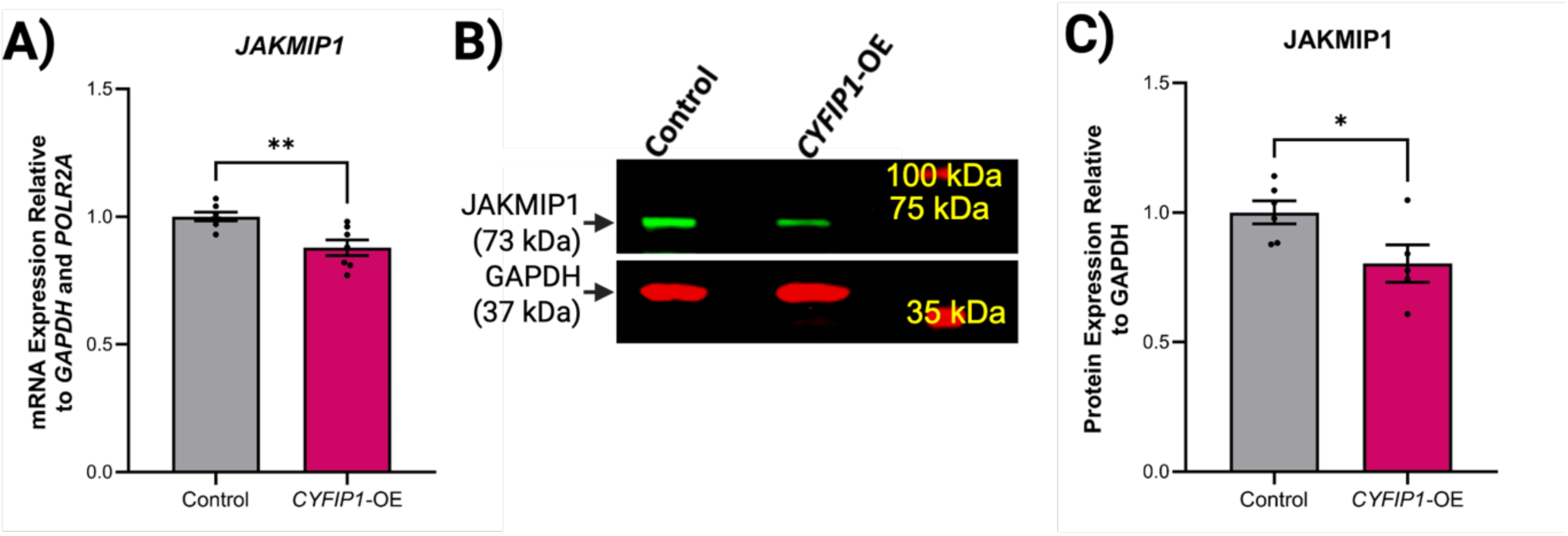
| CYFIP1-OE leads to reduced JAKMIP1 mRNA and protein expression. **A)** Relative mRNA expression of JAKMIP1 in Control and CYFIP1-OE cells measured by qRT-PCR. **B)** Western blotting of JAKMIP1 expression in Control and CYFIP1-OE cells. **C)** Densitometric analysis of total JAKMIP1 protein expression relative to the GAPDH loading control. For all experiments, values are presented as mean ± SEM of N = 3 independent experiments, 2-3 technical replicates each. Statistical significance against Control was determined using Unpaired Student’s T-Tests; *P < 0.05; **P < 0.01.

### *CYFIP1*-OE alters the expression of several cytokine signalling-related transcription factors

We have previously demonstrated that Dup(15q) iPSCs and iNeurons with increased *CYFIP1* expression show reduced *STAT3* expression (6). Furthermore, we have demonstrated that *JAKMIP1*-knockout SH-SY5Y cells also show reduced *STAT3* expression and that JAKMIP1 deficiency may alter other cytokine signalling pathways too (6). Therefore, to determine whether *CYFIP1*-OE may also lead to altered expression of cytokine signalling-related transcription factors, qRT-PCR was performed to measure mRNA expression of *STAT1*, *STAT3*, *STAT5B* and *NFKB1*. We observed that the expression of *STAT1* (Figure 3A) and *STAT3* (Figure 3B) were reduced in *CYFIP1*-OE HEK293 cells, whereas the expression of *STAT5B* (Figure 5C) and *NFKB1* (Figure 5D) were increased. This suggests that CYFIP1 may also modulate the expression of cytokine signalling-related transcription factors, likely via modulation of *JAKMIP1*.

**Figure 3.**
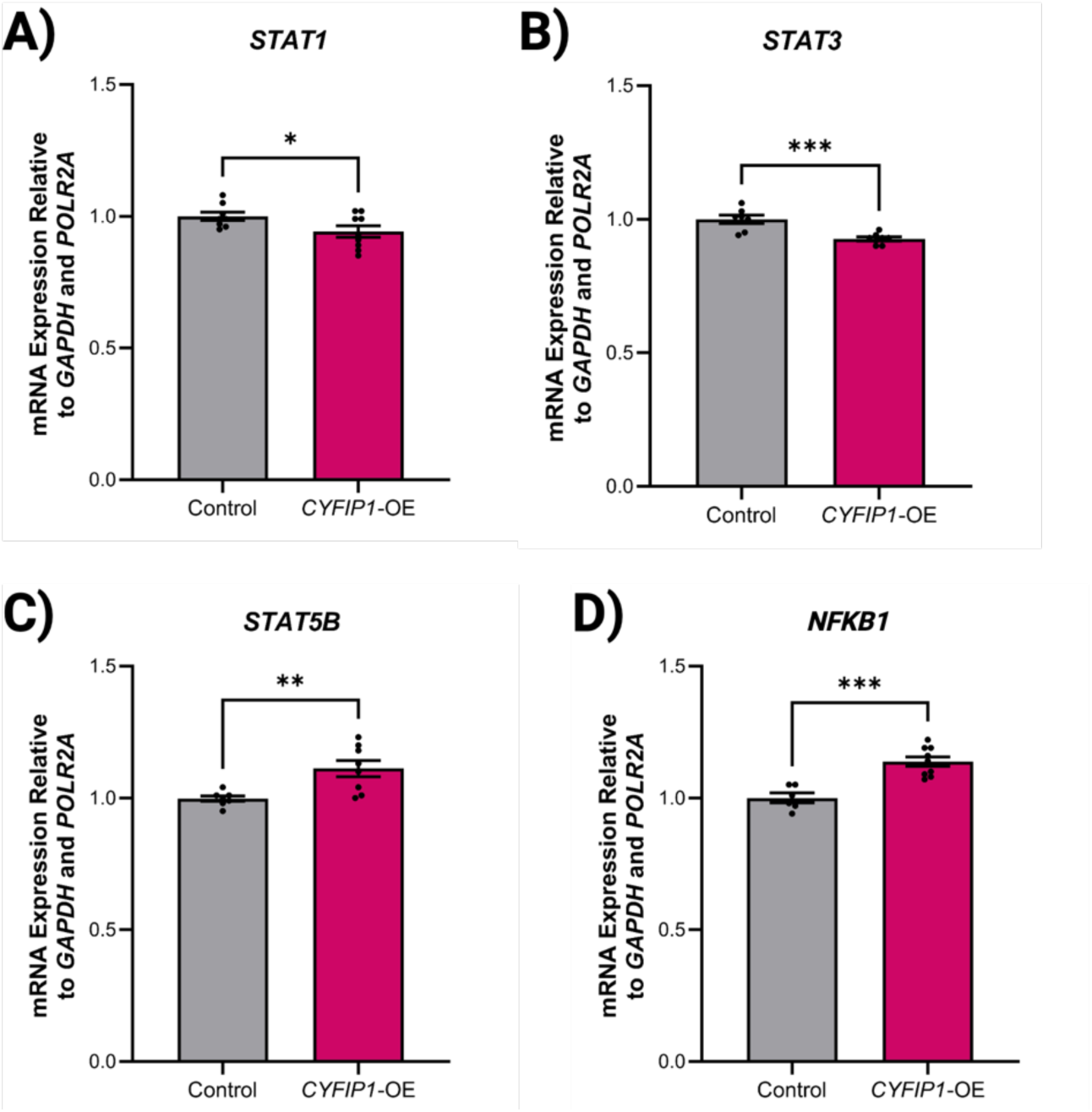
| CYFIP1-OE alters the expression of other cytokine signal-regulated transcription factors. Relative mRNA expression of **A)** STAT1, **B)** STAT3, **C)** STAT5B or **D)** NFKB1 in Control and CYFIP1-OE cells measured by qRT-PCR. For all experiments, values are presented as mean ± SEM of N = 3 independent experiments, 2-3 technical replicates each. Statistical significance against Control was determined using Unpaired Student’s T-Tests; *P < 0.05; **P <0.01; ****P < 0.0001.

**Figure 4.**
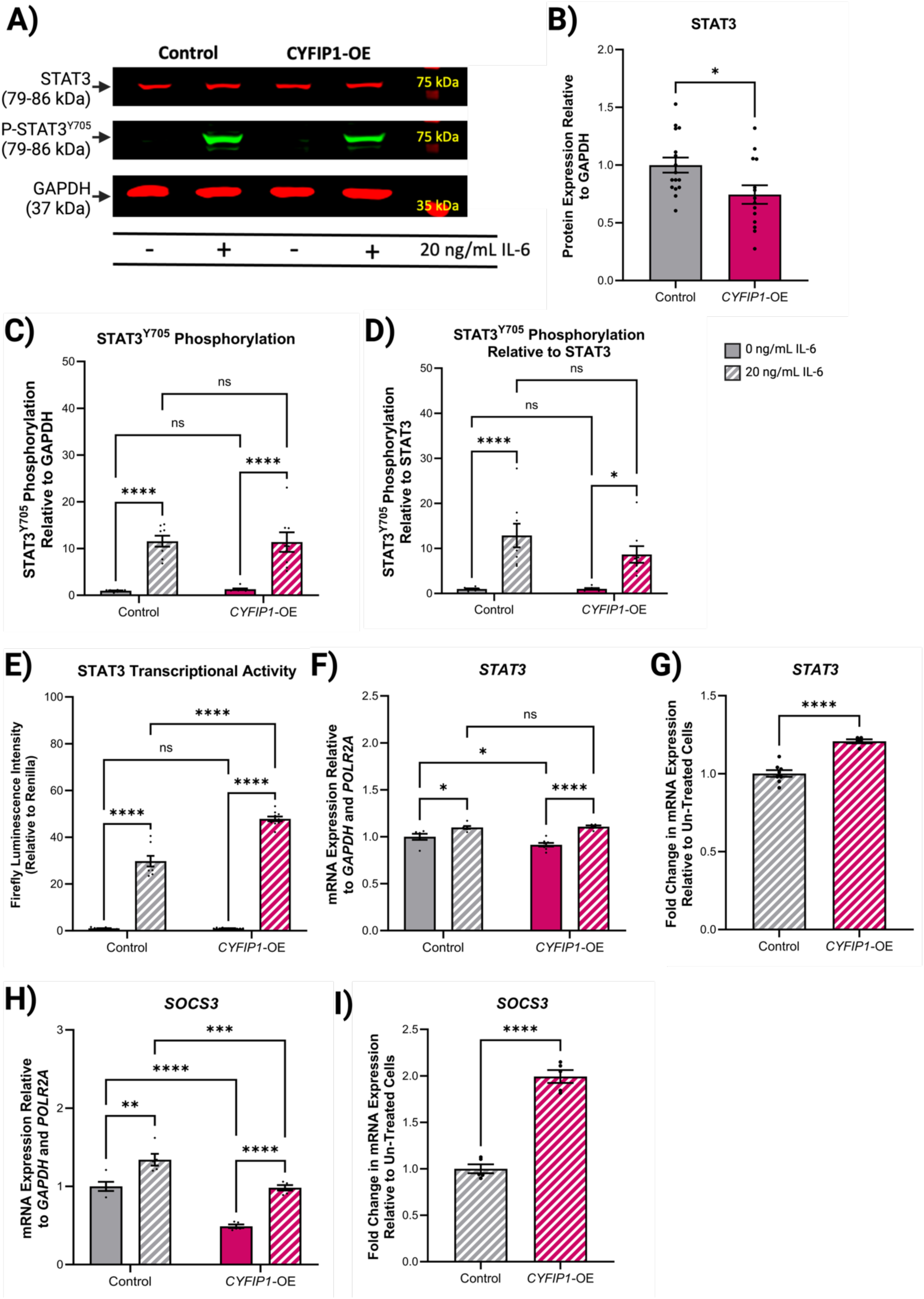
| CYFIP1-OE enhances STAT3 transcriptional activity but not STAT3 activation. **A)** Western blotting of STAT3 expression and phosphorylation (P-STAT3^Y705^) in Control and CYFIP1-OE cells following 30-minute treatment with 0 ng/mL IL-6 or 20 ng/mL IL-6. Densitometric analysis of **B)** total STAT3 protein expression relative to the GAPDH loading control, **C)** P-STAT3 relative to the GAPDH loading control or **D)** P-STAT3 relative to total STAT3 protein expression. **E)** Total transcriptional activity at a STAT3-regulated promoter in Control and CYFIP1-OE cells stimulated with or without 20 ng/mL IL-6 for 18 hours post-transfection with Cignal Reporter plasmids. Firefly luminescence intensity is expressed as a ratio to Renilla luminescence intensity to adjust for transfection emiciency. **F)** Relative mRNA expression of STAT3 in Control and CYFIP1-OE cells treated with 0 ng/mL or 20 ng/mL IL-6 for 4 hours, measured by qRT-PCR. **G)** Fold change in STAT3 mRNA expression following IL-6 treatment relative to untreated cells in part F). Values presented as mean ± SEM of N = 3 independent experiments, 3 technical replicates each. Statistical significance against Control was determined using Unpaired Student’s T-Tests; ****P < 0.0001. **H)** Relative mRNA expression of SOCS3 in Control and CYFIP1-OE cells treated with 0 ng/mL or 20 ng/mL IL-6 for 4 hours, measured by qRT-PCR. **I)** Fold change in SOCS3 mRNA expression following IL-6 treatment relative to untreated cells in part H). For all experiments, values are presented as mean ± SEM of N = 3 independent experiments, 3-4 technical replicates each. Statistical significance against Control was determined using Unpaired Student’s T-Tests when comparing two groups, or two-way ANOVAs with Tukey’s Honest Significant Dimerence tests when comparing multiple groups; ns – P > 0.05; *P < 0.05; **P < 0.01; ***P < 0.001; ****P < 0.0001.

**Figure 5.**
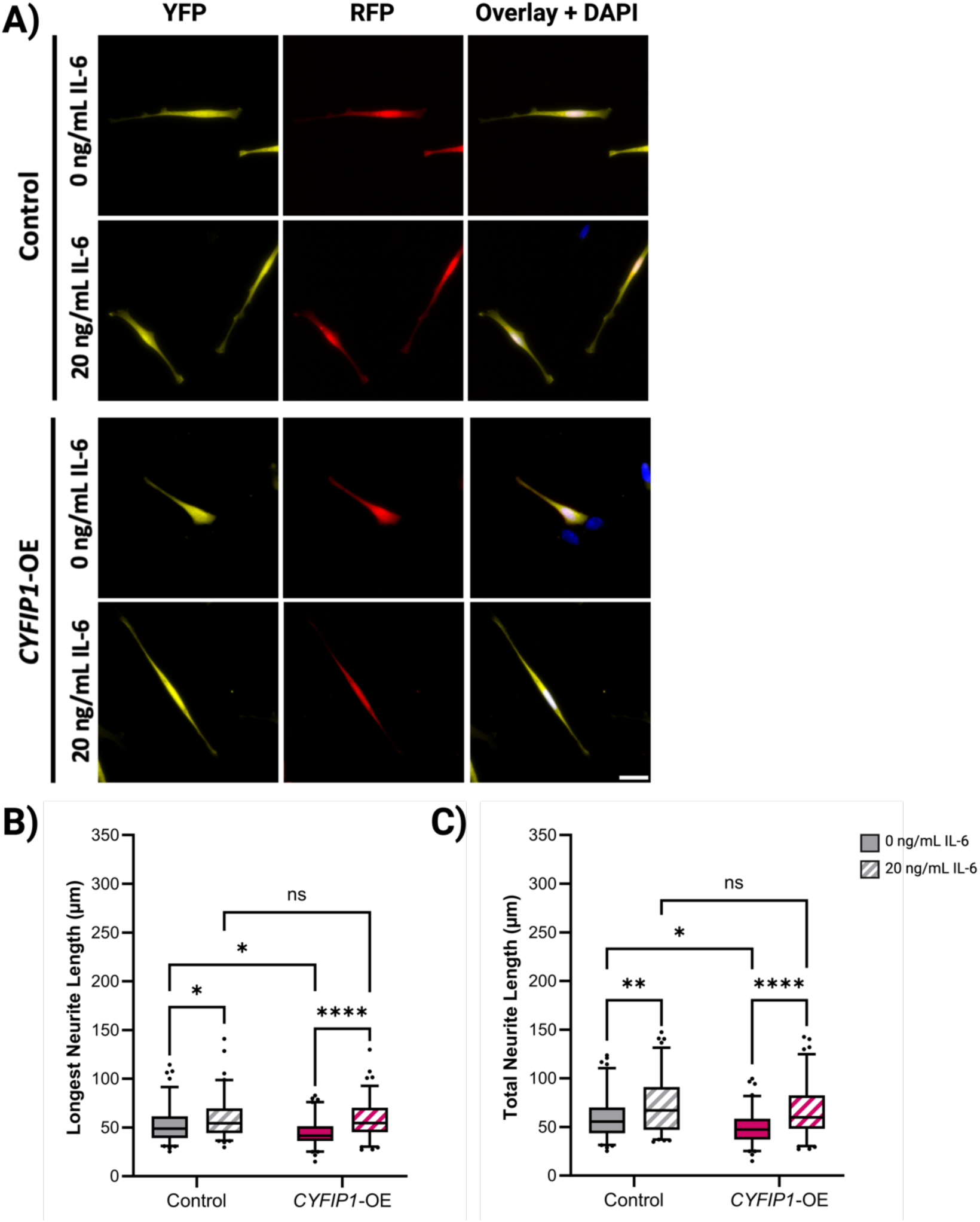
| CYFIP1-OE enhances IL-6-induced neuritogenesis. SH-SY5Y cells were Nucleofected with either the RFP-T2A backbone vector and pCAβ-YFP as the control condition; or RFP-T2A-CYFIP1 and pCAβ-YFP to assess the emects of CYFIP1-OE on IL-6-influenced neuritogenesis. Cells were dimerentiated for two days with 20 ng/mL BDNF and 10 μM RA under serum starvation with or without 20 ng/mL IL-6, fixed with 4% PFA and stained with DAPI. **A)** Fluorescence micrographs of dimerentiated SH-SY5Y cells. Images captured using an upright Leica DM4000 B LED microscope. Scale bar = 50 μm. Neurite tracing analysis was performed using the FIJI software. **B)** The length of the longest neurite produced by the cell was measured for all conditions. **C)** The sum length of all neurites produced per cell is presented for each condition. For all experiments, values are presented as mean ± SEM of N = 4 independent experiments, 3 technical replicates each. Statistical significance against Control was determined using two-way ANOVAs with Tukey’s Honest Significant Dimerence tests; ns – P > 0.05; *P < 0.05; **P < 0.01; ****P < 0.0001.

### *CYFIP1*-OE enhances IL-6-induced STAT3 activity

To investigate whether reduced *STAT3* expression in *CYFIP1*-OE HEK293 cells leads to altered cytokine-induced STAT3 responses we first performed Western blotting to assess IL-6-induced STAT3^Y705^ phosphorylation (representative of STAT3 activation) (Figure 4A). Despite the reduction in STAT3 expression (Figure 4B), *CYFIP1*-OE cells still show strong responses to IL-6, demonstrating no significant difference in IL-6-stimulated STAT3^Y705^ phosphorylation levels compared to control cells, both at the ‘whole-cell level’ (when STAT3^Y705^ phosphorylation levels are normalised to GAPDH) (Figure 4C) and upon closer examination of the phosphorylation kinetics of STAT3 itself (when STAT3^Y750^ phosphorylation levels are normalised to total STAT3 protein levels) (Figure 4D). This suggests that although *CYFIP1*-OE cells show reduced STAT3 expression, STAT3 activation following IL-6 stimulation is unaltered.

STAT3 activity is modulated by various post-translational modifications in addition to phosphorylation at the Y705 residue. Therefore, despite observing no change in STAT3^Y705^ phosphorylation following *CYFIP1*-OE, we examined whether *CYFIP1*-OE cells show altered IL-6-induced transcriptional activity by performing a DLR™ assay. This assay places the control of a Firefly luciferase under a STAT3 response element, meaning that increased Firefly luminescence is indicative of increased STAT3 transcriptional activity. Using this DLR™ assay, we found that *CYFIP1*-OE cells show increased Firefly luminescence following IL-6 stimulation than control cells (Figure 4E), suggesting that IL-6-induced STAT3 transcriptional activity is enhanced in *CYFIP1*-OE cells regardless of the reduced STAT3 expression.

To validate the DLR™ assay results, qRT-PCR was used to measure the mRNA expression of STAT3-responsive genes in *CYFIP1*-OE cells following IL-6 stimulation. Somewhat contradictory to the DLR™ assay, levels of *STAT3* mRNA in IL-6-treated *CYFIP1*-OE cells were not higher than those in untreated control cells (Figure 4F). However, considering that *CYFIP1*-OE cells already show reduced *STAT3* mRNA expression at baseline, this is unsurprising. Therefore, after normalising the levels of IL-6-stimulated *STAT3* expression to unstimulated *STAT3* expression, *CYFIP1*-OE cells show an enhancement in IL-6-induced *STAT3* mRNA expression compared to control cells (Figure 4G). The expression of *SOCS3* shows a similar phenomenon whereby IL-6-stimulated *CYFIP1*-OE cells appear to show lower *SOCS3* mRNA levels compared to IL-6-treated control cells (Figure 4H), however the fold change in *SOCS3* expression relative to unstimulated cells shows that IL-6-induced *SOCS3* expression is magnified in *CYFIP1*-OE cells (Figure 4I). Taken together, these findings complement the DLR™ assay results, suggesting that *CYFIP1*-OE cells show enhanced STAT3 transcriptional activity following IL-6 stimulation.

### *CYFIP1*-OE enhances IL-6-induced neuritogenesis

To understand how this enhanced STAT3 response to IL-6 in *CYFIP1*-OE cells observed at a molecular level affects a broader IL-6-stimulated cellular process, we differentiated *CYFIP1*-OE SH-SY5Y cells in the presence of IL-6 and measured the neurite lengths produced (Figure 5A), as we have previously shown that IL-6 induces neurite outgrowth in SH-SY5Y cells (6). As expected, control and *CYFIP1*-OE cells differentiated in the presence of IL-6 exhibit longer LNL (Figure 5B) and TNL (Figure 5C) measurements than cells differentiated without IL-6. *CYFIP1*-OE cells appear to have enhanced neurite outgrowth responses to IL-6, as although *CYFIP1*-OE cells produce shorter neurites compared to the control as baseline, the LNL (Figure 5B) and TNL (Figure 5C) measurements are no longer significantly different between the two populations when differentiated with IL-6. These data appear to agree with the suggestion that *CYFIP1*-OE leads to enhanced STAT3 responses to IL-6, and that this enhancement in IL-6-induced STAT3 activity leads to increased IL-6-stimulated neurite extension and production.

### *CYFIP1*-OE enhances IFN-γ-induced STAT1 transcriptional activity

Having determined that *CYFIP1*-OE alters the expression of several cytokine signalling-related transcription factors, and that *CYFIP1*-OE enhances IL-6-induced STAT3 activity, we aimed to investigate whether *CYFIP1*-OE alters the activity of another cytokine signalling-related transcription factor. STAT1 was chosen to follow up with because similar to *STAT3* expression, the expression of *STAT1* was also reduced in *CYFIP1*-OE HEK293 cells. A DLR™ assay which placed the control of a Firefly luciferase under a GAS element (the response element STAT1 binds to initiate transcription) was performed. We found that *CYFIP1*-OE HEK293 cells show increased Firefly luminescence following IFN-γ stimulation compared to control cells (Figure 6A), suggesting that IFN-γ-induced STAT1 transcriptional activity is enhanced in *CYFIP1*-OE cells, again regardless of the reduced *STAT1* expression.

**Figure 6.**
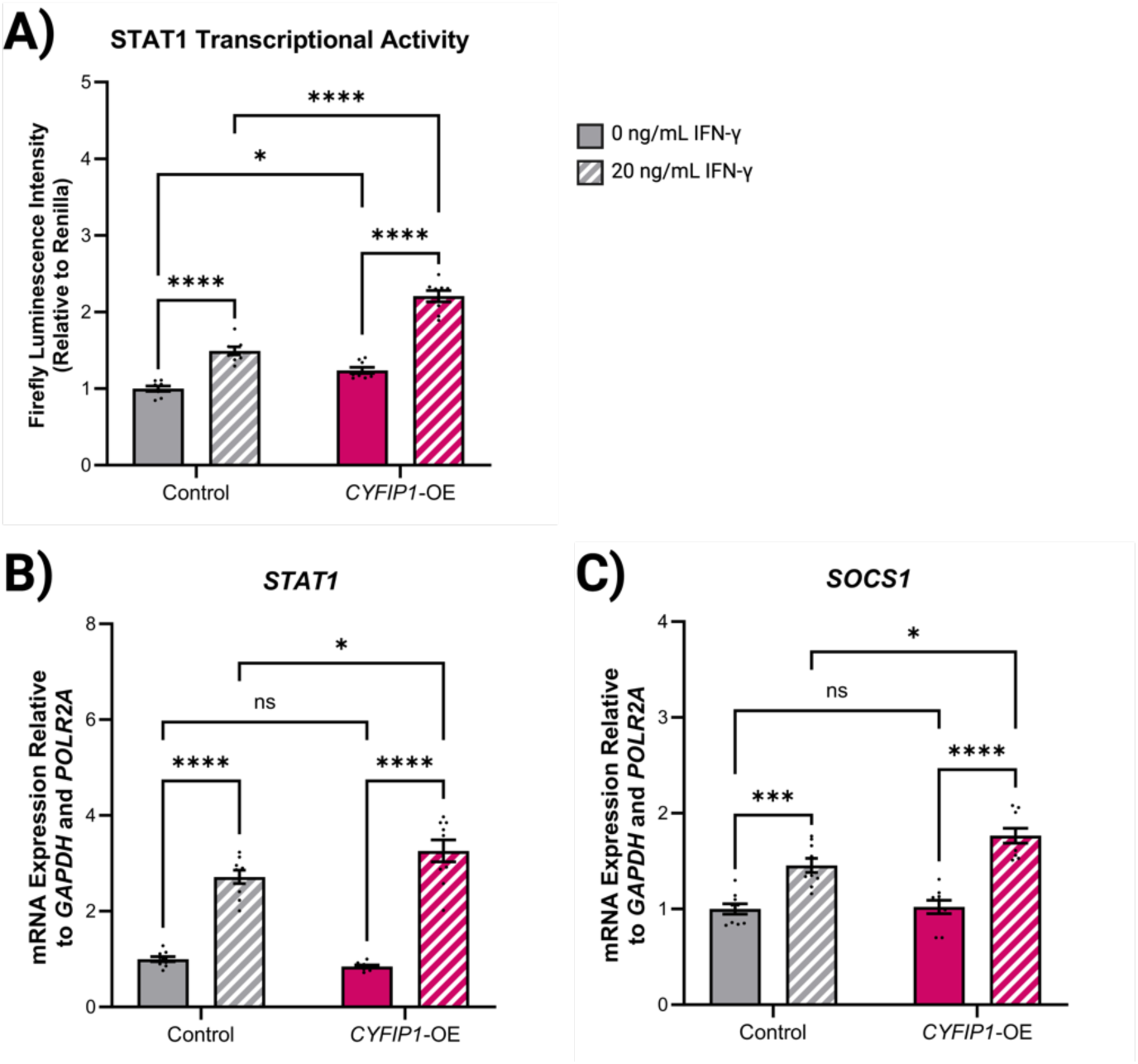
| CYFIP1-OE leads to enhanced STAT1 transcriptional activity following IFN-γ treatment. **A)** Total transcriptional activity at a STAT1-regulated promoter in Control and CYFIP1-OE cells stimulated with or without 20 ng/mL IFN-γ for 18 hours post-transfection with Cignal Reporter plasmids. Firefly luminescence intensity is expressed as a ratio to Renilla luminescence intensity to adjust for transfection emiciency. **B)** Relative mRNA expression of STAT1 in Control and CYFIP1-OE cells treated with 0 ng/mL or 20 ng/mL IFN-γ for 4 hours, measured by qRT-PCR. **C)** Relative mRNA expression of SOCS1 in Control and CYFIP1-OE cells treated with 0 ng/mL or 20 ng/mL IFN-γ for 4 hours, measured by qRT-PCR. For all experiments, values are presented as mean ± SEM of N = 4 independent experiments, 3 technical replicates each. Statistical significance against Control was determined using two-way ANOVAs with Tukey’s Honest Significant Dimerence tests; ns – P > 0.05; *P < 0.05; ***P < 0.001; ****P < 0.0001.

As validation of the DLR™ assay, qRT-PCR was performed to measure the expression of STAT1-response genes following IFN-g stimulation. *CYFIP1*-OE cells show increased IFN-γ-induced *STAT1* expression (Figure 6B), suggesting an enhanced transcription response to IFN-γ stimulation. Additionally, although not significantly different at baseline, *CYFIP1*-OE cells show increased *SOCS1* expression following IFN-γ treatment (Figure 6C), again suggesting an enhanced STAT1 transcriptional response to IFN-γ stimulation. This also suggests that the observed alterations in IL-6/STAT3 signalling are not limited to STAT3 and that CYFIP1 may play a role in modulating a larger range of cytokine signalling pathways.

## Discussion

### A novel role for CYFIP1 in modulating IL-6/STAT3 signalling independent from JAKMIP1

Nishimura *et al*. (5) first linked *JAKMIP1* to ASD and Dup(15q), reporting that *JAKMIP1* expression was altered in Dup(15q) and that overexpression of the chromosome 15q gene *CYFIP1* correspondingly resulted in *JAKMIP1* downregulation. We expanded upon this in our previous work, identifying JAKMIP1 as a modulator of IL-6 cytokine signaling in SH-SY5Y cells, and demonstrating that Dup(15q) iPSCs and cortical neurons with elevated *CYFIP1* dosage and expression exhibit enhanced IL-6-induced STAT3 activity (6). However, given that the magnified IL-6 signaling observed in these Dup(15q) cells did not correlate with *JAKMIP1* expression levels, we hypothesised that another gene within the 15q region, likely *CYFIP1*, might be driving this increased responsiveness to IL-6.

To test this hypothesis, we induced *CYFIP1* overexpression in HEK293 cells and discovered that this resulted in the altered expression of several cytokine signaling-related transcription factors. In particular, *STAT1* and *STAT3* expression were reduced, whereas *STAT5B* and *NFKB1* expression were raised, suggesting that CYFIP1 may potentially exert broad cellular functions that impact the signaling networks of multiple cytokines. Taking this forward, we measured IL-6-induced activation and activity of STAT3. Consistent with the Dup(15q) iPSCs and neurons from our previous work, we observed that despite the reduced *STAT3* expression, IL-6-induced STAT3 responses were enhanced in *CYFIP1*-OE HEK293 cells. Furthermore, *CYFIP1* overexpression also potentiated IL-6-triggered neuritogenesis in SH-SY5Y cells. As such, this study provides the first evidence that CYFIP1 may play a role in regulating IL-6/STAT3 signalling in both HEK293 and SH-SY5Y cells, and its upregulation could be responsible for the enhanced responsiveness to IL-6 exhibited by Dup(15q) iPSCs and neurons.

### How does CYFIP1 influence STAT1 and STAT3 activity?

The mechanisms underlying CYFIP1-mediated regulation of *STAT1* and *STAT3* expression and activity remain unclear. Regarding STAT3, reduced *STAT3* expression in *JAKMIP1*-deficient cells (6) may suggest that JAKMIP1, rather than CYFIP1, is responsible for regulating *STAT3* expression. This could occur through its interactions with various RNA-binding proteins (RBPs) or splicing factors (6). However, it is important to note that *CYFIP1* overexpression enhances IL-6-induced STAT3 transcriptional activity, whereas *JAKMIP1*-knockout impairs it (6). Thus, whilst JAKMIP1 could be responsible for controlling *STAT3* expression, when considering STAT3 activity, our results indicate that it is more likely that CYFIP1 is responsible by a mechanism independent of JAKMIP1.

Interestingly, despite *CYFIP1* overexpression leading to increased STAT3 transcriptional activity following IL-6 treatment, we observed no significant difference in the levels STAT3^Y705^ phosphorylation in *CYFIP1*-OE cells, suggesting that CYFIP1 does not alter STAT3 phosphorylation kinetics, and thus STAT3 activation. One reason for this discrepancy could be because of the influence of other post-translational modifications of STAT3. For example, STAT3^K685^ acetylation may be necessary for IL-6-induced STAT3-responsive gene expression (16). Rather than modulating STAT3 activation through phosphorylation, perhaps CYFIP1 may somehow control STAT3 activation through acetylation, however, the exact mechanism by which CYFIP1 could regulate acetylation is unclear.

In terms of *STAT1* expression and activity, it would not be unreasonable to suggest that these changes in STAT1 activity may be mediated by the changes in *JAKMIP1* expression. JAKMIP1 was previously reported to inhibit Interferon-α (IFN-α) signalling (17). Importantly, both IFN-α and IFN-γ signalling cascades involve the phosphorylation and activation of STAT1. It is therefore likely that CYFIP1-induced downregulation of *JAKMIP1* may reduce the inhibiting influence of JAKMIP1 on IFN signalling, thereby enhancing the IFN response. Furthermore, STAT1 activity is also known to be altered by acetylation, similar to STAT3 (18). If CYFIP1-mediated regulation of STAT3 is dependent on post-translational modifications, the same mechanism(s) could be responsible for CYFIP1 regulation both STAT3 and STAT1 activity. Of course, all of these possibilities would require further validation.

### Relevance to ASD and neurodevelopment

Cytokine signalling plays crucial roles in neurodevelopment beyond immune responses, with STAT3 and STAT1 signalling in particular regulating several key processes. For example, STAT3 signalling has been demonstrated to control the switch between neurogenesis and gliogenesis, promoting both neural progenitor cell self-renewal and astrocyte differentiation (19). Similarly, STAT1 signalling has been shown to be involved in neurogenesis, and promoting neurite outgrowth (20). Given how crucial STAT3 and STAT1 signalling are for neurodevelopment, it is unsurprising that alterations in STAT3 and STAT1 signalling are associated with NDDs such as ASD (21,22). MIA, where an immune response is triggered by infection of exposure to immunogenic material during pregnancy, is associated with increased incidence of ASD (23). Both IL-6 and IFN-γ are pro-inflammatory cytokines that have been implicated in the pathogenesis of ASD, with IL-6 being a key player in MIA-induced ASD-like behaviours (23,24). Considering that *CYFIP1*-OE enhances STAT3 and STAT1 responses to IL-6 and IFN-γ respectively suggests that CYFIP1 may play a pivotal role in amplifying the inflammatory signals during MIA, potentially exacerbating the effects of immune activation on foetal brain development.

Amplification of both the IL-6/STAT3 and IFN-γ/STAT1 signalling pathways by CYFIP1 provides a potential molecular mechanism that explains the overlap between immune activation and ASD, particularly in cases where no obvious inflammatory trigger is identified. Elevated cytokine levels are frequently reported in individuals with ASD (25), and our results suggest that *CYFIP1* overexpression could act as a molecular convergence point, increasing the sensitivity of the developing brain to immune signals even under baseline inflammatory conditions. This could help to explain cases of ASD where immune activation is not readily apparent but inflammation-induced neurodevelopmental disruption still occurs.

## Conclusion

In summary, this study identifies a novel role for CYFIP1 in regulating the expression of cytokine signalling-related transcription factors, as well as enhancing their activity following cytokine stimulation. By amplifying these inflammatory pathways, CYFIP1 could play a central role in exacerbating the neuroinflammatory environment, leading to altered neurodevelopment and increased susceptibility to ASD. Further research is required to understand the mechanism by which CYFIP1 modulates these cytokine signalling pathways and to investigate this in the context of ASD with MIA.

## List of abbreviations

ASD: Autism spectrum disorder
Dup(15q): Chromosome 15q-duplication syndrome
CYFIP1: Cytoplasmic FMR1-intercating protein 1
iPSC: Induced pluripotent stem cell
IL-6: Interleukin-6
STAT3: Signal transducer and activator of transcription 3
OE: Overexpression
MIA: Maternal immune activation
JAKMIP1: Janus kinase and microtubule-interacting protein 1
FXS: Fragile X syndrome
iNeurons: iPSC-derived neurons
FMRP: Fragile X messenger ribonuclear protein
WRC: WAVE regulatory complex
SMAD3: SMAD family member 3
NF-kB: Nuclear factor of κB
ISRE: Interferon response element
TGF-β1: Transforming growth factor β1
GAPDH: Glyceraldehyde-3-phosphate dehydrogenase
DLR™: Dual-luciferase reporter™
SOCS3: Suppressor of cytokine signalling 3
LNL: Longest neurite length
TNL: Total neurite length
IFN-γ: Interferon-γ
RBP: RNA-binding protein

## Declarations

### Ethics approval and consent to participate

Not applicable.

### Consent for publication

Not applicable.

### Availability of data and materials

The datasets used and/or analysed during the current study are available from the corresponding author on reasonable request.

### Author’s contributions

E-R.M. conceived and designed the research, performed the laboratory research, generated the data and figures, and contributed to manuscript writing and editing. J.G.M. provided critical input and contributed to manuscript editing. M.A.R. provided critical input and reviewed the manuscript. A.O-A. conceived and designed the research, provided critical input, reviewed the manuscript, and secured funding. All authors reviewed and approved the final manuscript.

## Acknowledgements

The research was carried out at the National Institute for Health and Care Research (NIHR) Exeter Biomedical Research Centre (BRC). We gratefully acknowledge the generous support of the charitable organisations that contributed to this work, including the Wellcome Trust Translational Research Exchange @ Exeter (TREE) pump-priming fund and the Bristol Japanese Culture Showcase. We thank Liming Yang for valuable discussions during lab meetings, and we are also grateful to Dr. Kaiyven Afi Leslie for his technical advice and support.

## Supplementary Files

**Supplementary Figure 1.**
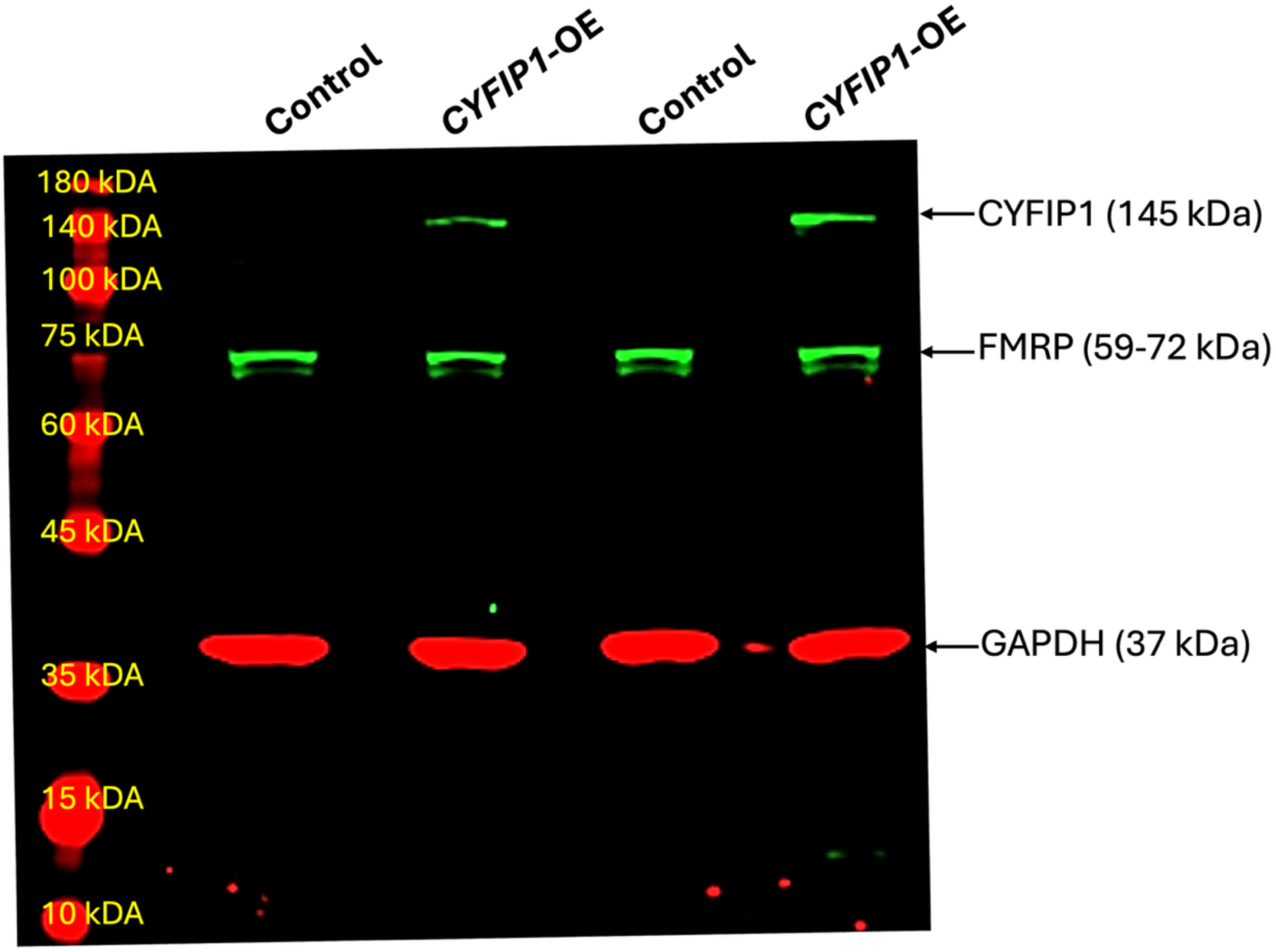
| Modelling CYFIP1 overexpression in HEK293 cells. Increased CYFIP1 protein expression in CYFIP1-OE cells 72 hours post-transfection confirmed via Western blotting. Membrane probed with: Rabbit Anti-CYFIP1 (#ab156016, Abcam), Rabbit Anti-FMRP (#13755-1-AP, Proteintech), Mouse Anti-GAPDH (#sc-47724, Santa Cruz Biotechnology), Goat Anti-Mouse DyLight™-680 (#35519, Invitrogen) and Goat Anti-Rabbit DyLight™-800 (#SA5-10036, Invitrogen).

**Supplementary Figure 2.**
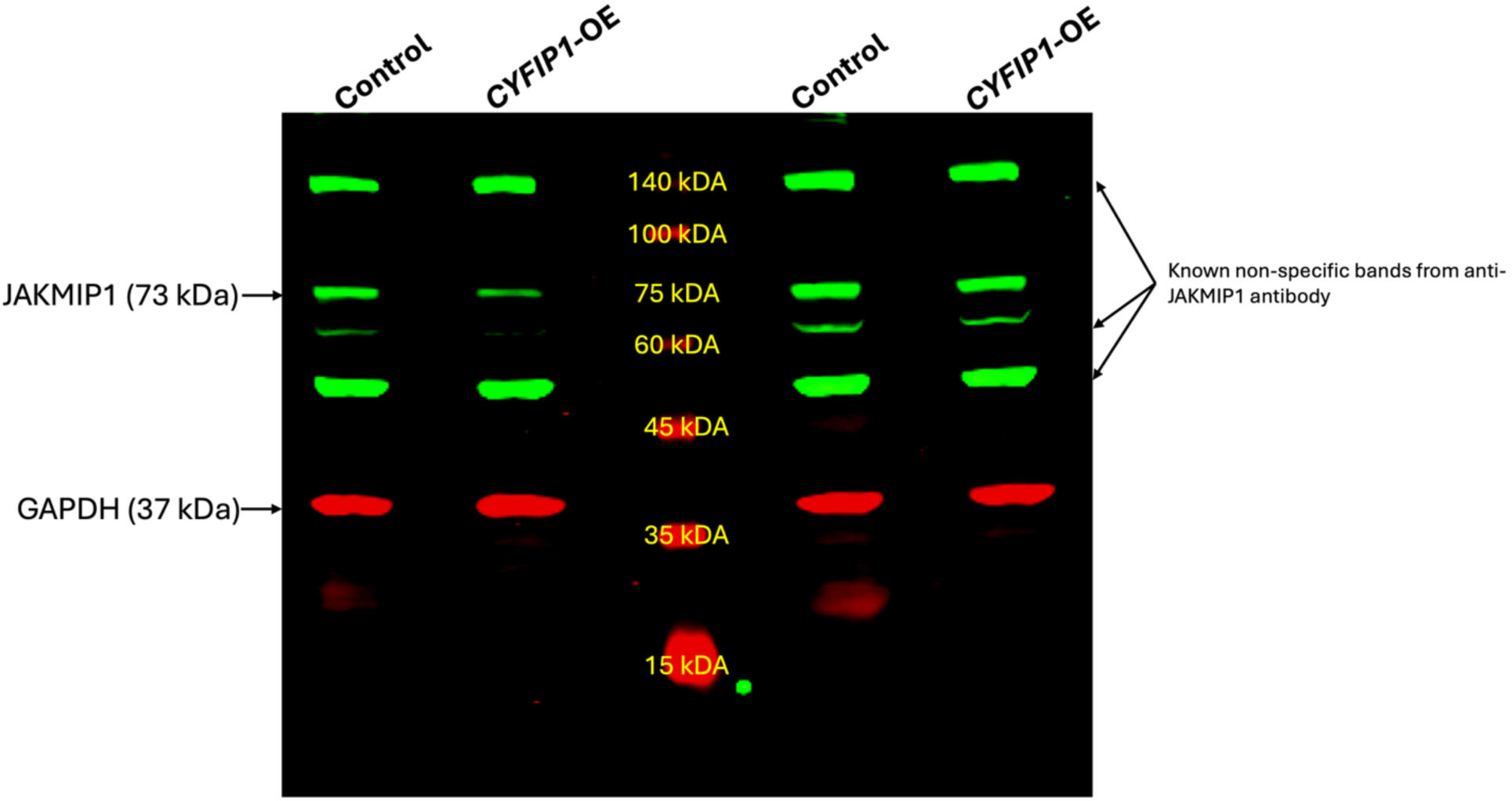
| CYFIP1-OE leads to reduced JAKMIP1 protein expression. Western blotting of JAKMIP1 expression in Control and CYFIP1-OE cells. Membrane probed with: Rabbit Anti-JAKMIP1 (#13846-1-AP, Proteintech), Mouse Anti-GAPDH (#sc-47724, Santa Cruz Biotechnology), Goat Anti-Mouse DyLight™-680 (#35519, Invitrogen) and Goat Anti-Rabbit DyLight™-800 (#SA5-10036, Invitrogen). Known non-specific bands produced are indicated on the blot.

**Supplementary Figure 3.**
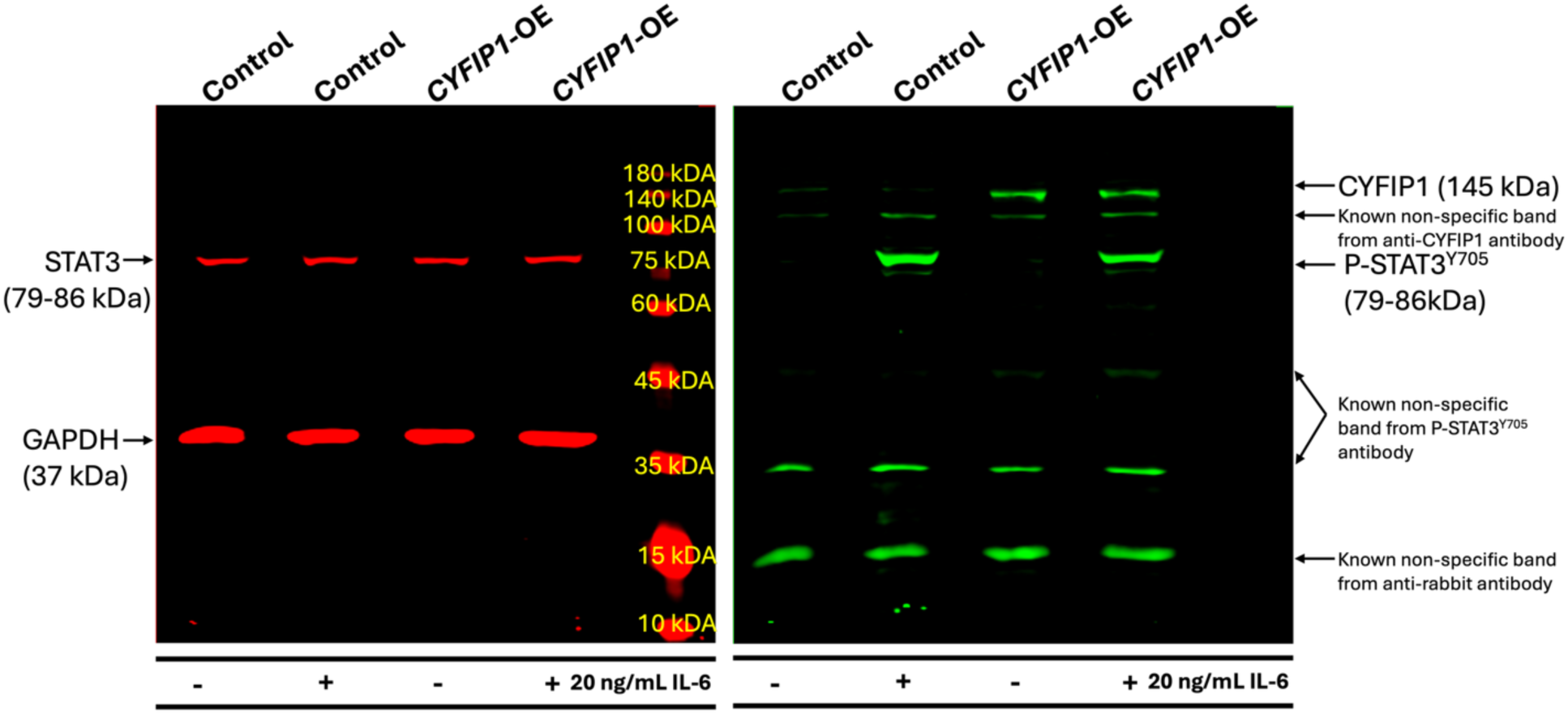
| CYFIP1-OE has no eYect on STAT3 activation. Western blotting of STAT3 expression and phosphorylation (P-STAT3^Y705^) in Control and CYFIP1-OE cells following 30-minute treatment with 0 ng/mL IL-6 or 20 ng/mL IL-6. Membrane probed with: Mouse Anti-STAT3 (#9139S, Cell Signaling Technology), Rabbit Anti-P-STAT3^Y705^ (#9131l, Cell Signaling Technology), Mouse Anti-GAPDH (#sc-47724, Santa Cruz Biotechnology), Goat Anti-Mouse DyLight™-680 (#35519, Invitrogen) and Goat Anti-Rabbit DyLight™-800 (#SA5-10036, Invitrogen). Known non-specific bands are indicated on the blot.

